# CD200R1 is required for the development of γδ17 T cells

**DOI:** 10.1101/2025.05.19.654867

**Authors:** Holly Linley, Shafqat Jaigirdar, Lucy Buckingham, Joshua Cox, Megan Priestley, Anna Hains, Amy Saunders

## Abstract

γδ T cells are enriched at barrier sites such as skin, gut and lung, where they protect against cancer and infections and promote healing. They detect diverse ligands in T cell receptor-dependent or independent manners, producing large quantities of pro-inflammatory cytokines. γδ T cells develop in foetal thymi in temporally controlled waves where, unlike αβ T cells, many γδ T cells adopt their effector fate, becoming either IFNγ or IL-17A-producers (γδ17 T cells).

CD200R1 suppresses myeloid cell activity but has also been shown to promote innate lymphoid cell IL-17A production, enhancing psoriasis-like skin inflammation. γδ17 T cells are potent IL-17A producers in skin therefore, the effect of CD200R1 on IL-17A production by γδ17 T cells was investigated. CD200R1 was found to promote IL-17A production by γδ T cells by supporting the development of γδ17 T cells, enhancing IL-17-producing and RORγt^+^ γδ T cell numbers in foetal thymic organ cultures. To fulfil this role, CD200R1 acts either directly on developing γδ T cells, or indirectly on thymic stromal cells. This identifies CD200R1 as a critical novel regulator of γδ17 T cell development in early life, a key process for ensuring immunity, particularly at barrier sites.

## Introduction

γδ T cells are rare in many tissues but are enriched at barrier sites such as the skin, lung and gut where they are crucial for wound healing and protecting against infections and cancer (1–5). However, they have also been shown to drive inflammatory disease (6). γδ T cells are potent and rapid producers of inflammatory cytokines, being activated in either a T cell receptor (TCR)-dependent, or independent manner. Like conventional αβ T cells, γδ T cells rearrange their TCR genes in the thymus however, γδ TCRs are less variable than αβ TCRs (5) and lack MHC restriction. Instead, γδ TCRs recognise a range of ligands including stress-induced MHC-like molecules, phosphorylated metabolites and lipid antigens presented by CD1 molecules (7).

γδ T cells can be classified into subtypes based on their effector capabilities, being either IFNγ or IL-17A-producers, with effector phenotypes conferred by the lineage-defining transcription factors, T-bet or RORγt respectively. Concomitantly with expressing rearranged γδ TCR chains, γδ T cells also acquire their effector fate, with murine Vγ1 and Vγ5 cells (Tonegawa nomenclature (8)) mainly gaining IFNγ-producing potential, and Vγ4 and Vγ6 T cells largely gaining IL-17A-producing potential (9). The Vγ TCR chain expressed is not the determining factor in effector fate acquisition (10, 11), but Vγ gene order on the TCRγ locus is important (12). The development of γδ T cells occurs in ordered waves comprising different subsets, beginning at embryonic day 15 of gestation with the appearance of Vγ5 (Dendritic epidermal T cells (DETC)) which home to the epidermis (13). This is followed by the development of Vγ6 cells which home to dermis, uterus and the peritoneal cavity. Vγ4 cells develop shortly after, and home to lung, dermis and lymph node (LN) (14), then perinatally, Vγ7 (intra-epithelial lymphocytes (IEL)) develop and home to the intestine. Perinatal progenitors are required for generating IL-17-producing γδ (γδ17) T cells in the absence of inflammation (11), however, γδ17 T cells can be generated *de novo* from adult bone marrow-derived precursors in response to inflammation (15) and Vγ4 and Vγ1 T cells continue to develop later in life but are ‘adaptive γδ T cells’ retaining the ability to adopt multiple effector fates in the periphery (16). Factors governing the sequential development of γδ T cell subsets are not completely understood, but both TCR and environmental signals are important.

TCR signal strength plays a crucial role in γδ T cell development. CD4^-^ CD8^-^ double negative (DN) cells receiving strong TCR signals commit to the γδ lineage and strong TCR signalling is key for IFNγ-producing γδ T cell development (17, 18). However, weaker TCR signals are required for γδ17 T cell development (14, 17–20), where distinct TCR-signalling pathways are engaged, specifying γδ17 T cell fate (21). Negative regulators of TCR signalling such as the Src family kinase Blk, aid the development of γδ17 T cells (22), as does the transcription factor cMaf which drives expression of the lineage defining transcription factor, *Rorc.* TCR signal strength regulates cMaf levels, with strong TCR signals reducing cMaf, and weaker TCR signals increasing cMaf levels and thus driving *Rorc* and reinforcing γδ17 T cell effector fate (23).

Cytokines also regulate γδ T cell development with IL-7Rα required for V-J γδTCR recombination (24–28). IL-7 signalling particularly impacts γδ17 T cells, as high levels of IL-7/IL-7Rα signalling favour IL-17 production (29, 30) and IL-7Rα is required for the homeostasis of γδ17 T cells (31). Lymphotoxin (LT) signalling is key for the development of both IFNγ-producing and γδ17 T cells, where LTβR drives the NF-κB family member, RelB, which in turn maintains *Rorc* expression (32). TGFβ also drives γδ T cell IL-17 production in early life (33), although the mechanism is not yet understood.

Medullary thymic epithelial cells (mTECs) and cortical TECs (cTECs) are crucial for conventional αβ T cell development, supporting positive and negative selection and providing Notch signals and IL-7 (34). However, less is known about their role in supporting γδ T cell development. cTECs promote Vγ4 T cell activity but are not required for Vγ6 T cells (35). Conversely, mTECs are important for Vγ5 DETC development, as they provide signals via the butyrophilin-related protein, Skint-1 (36). Thymic dendritic cells (DCs) also contribute to conventional αβ T cell development, but a role in γδ T cell development has not been defined. Despite advances in our understanding of how γδT cell development is regulated, factors involved in this process remain incompletely understood.

CD200R1 is a cell surface receptor expressed on many immune cell types, inhibiting pro-inflammatory cytokine production and mast cell degranulation in myeloid cells (37, 38). However, CD200R1 is required for efficient IL-17 production by innate lymphoid cells (ILCs), and CD200R1 promotes inflammation in a psoriasis model and controls cutaneous fungal infections (39, 40), demonstrating that CD200R1 has diverse functions in different cell types, including promoting IL-17 production.

Given that γδ T cells are crucial for producing IL-17 in skin and drive psoriasis-like and anti-fungal immune responses (41–43), the role of CD200R1 in regulating γδ T cell IL-17 production was investigated. We show that CD200R1-deficient mice have impaired IL-17 production by γδ T cells, due to a specific reduction in γδ17 T cells, with IFNγ-producing cells not affected. The reduction in γδ17 T cells is due to defects in development, with CD200R1 likely required on either the developing γδ T cells themselves, or on thymic dendritic cells. Therefore, here we identify for the first time CD200R1 as a key factor required for the development of γδ17 T cells, shaping immunity at barrier sites, but also potentially contributing to the inflammatory skin disease psoriasis.

## Materials and Methods

### Mice

All animal experiments were locally ethically approved and performed in accordance with the UK Home Office Animals (Scientific Procedures) Act 1986. C57BL/6 (WT) mice were obtained from Charles River Laboratories. CD200R1KO mice (44) on a C57BL/6 background, were bred and maintained in specific pathogen free conditions in house. Mice were male and 7 to 12 weeks old at the start of procedures, unless otherwise stated. Where neonatal and foetal thymi were used, sex was not determined.

### Skin cell isolation

Ears were split in half and floated on 0.8% w/v Trypsin (Sigma) at 37 °C for 30 min, then were chopped and digested in 0.1 mg/ml (0.5 Wunch units/ml) Liberase TM (Roche) at 37 °C for 1 hr. The fat was removed from dorsal skin before floating on Trypsin as above. Tissue was chopped and digested with 1 mg/ml Dispase II (Roche) at 37 °C for 1 hr. Digested skin tissue suspensions were passed through 70 µm cell strainers, washed and counted.

### Lymph node, thymus and spleen cell isolation

Lymph nodes (LN, inguinal, axillary and brachial), thymi or spleens were passed through 70 µm cell strainers and washed. Red blood cells present in the splenocyte samples were lysed on ice for 3 min with ACK lysing buffer (Lonza), before cells were washed and counted.

### Flow cytometric analysis of cells

Cells were incubated with 0.5 µg/ml anti-CD16/32 (2.4G2, BD Bioscience) and Near IR Dead cell stain (Invitrogen) (except when Annexin V/7AAD staining was performed) prior to staining with fluorescently labelled antibodies. Cells were fixed with Foxp3/Transcription Factor Buffer Staining Set (eBioscience) for between 30 min and 16 hr at 4°C.

For cytokine analysis, 10 μM Brefeldin A was added to cell cultures for 4 hr prior to staining for cell surface markers as described above. After overnight fixation, cells were permeabilized with Foxp3/Transcription Factor Buffer Staining Set (eBioscience) and were stained with antibodies against intracellular markers or cytokines. Cells were analysed on a Fortessa, LSRII (both BD Bioscience) or Cytoflex (Beckman) flow cytometer. Data were analysed using FlowJo (TreeStar). Antibodies are detailed in Supplementary Table 1.

For Annexin V/7AAD staining, cells were stained on ice for cell surface markers, then were stained with Annexin V (Biolegend) following the manufacturer’s instructions. 7AAD (Biolegend) was added and cells were analysed within 2 hours.

For Vγ6 staining, cells were stained with 17D1 hybridoma supernatant for 1 hr, then were incubated with Biotinylated anti-Rat IgM (Sigma), followed by PECF594-conjugated streptavidin (eBioscience), each for 30 min, before being stained with other antibodies to detect cell surface markers.

### *In vitro* γδ T cell activations

Mouse dorsal skin cells were cultured in complete RPMI (RPMI supplemented with 10% heat inactivated FBS, 1% penicillin streptomycin solution, 2 mM L-glutamine, 1 mM sodium pyruvate, 20 mM HEPES, 1X non-essential amino acid solution, 25 nM 2-mercaptoethanol (Sigma)) and stimulated with 40 ng/mL IL-23 (Biolegend) for 16 hr. Cells were cultured with 10 μM Brefeldin A (Sigma) for 4 hr before staining for flow cytometric analysis. Alternatively, dorsal skin cells, thymocytes, splenocytes or LN cells were stimulated with 50 ng/mL PMA and 500 ng/mL ionomycin (Sigma) and 10 μM Brefeldin A for 4 hr before staining for flow cytometric analysis.

For co-culture of WT and CD200R1KO cells, the cells were isolated from dorsal skin then either the WT or CD200R1KO cells were labelled with 10 μM eF450-conjugated cell proliferation dye (Fischer Scientific) for 10 minutes in the dark at 37°C before washing and co-culturing with unlabelled cells.

### Foetal thymic organ cultures (FTOCs)

FTOCs were set up and analysed as described previously (14). Briefly, E15-15.5 embryos were obtained from WT or CD200R1KO timed matings. Foetal thymic lobes were dissected and cultured on nucleopore membrane filters (Whatman) (4-5 per filter) in complete RPMI (as above) for 8 days. Where necessary, cells were stimulated with 50 ng/ml PMA (Sigma) and 500 ng/ml Ionomycin (Sigma) with 10 μM Brefeldin A (Sigma) and 2 μM Monensin (eBioscience) for 4 hrs. Thymic lobes were homogenised using a syringe plunger and were stained and analysed by flow cytometry as described above. To count cells, Precision count beads (BioLegend) were added to each FTOC sample prior to analysis.

### Statistical analysis

Data were analysed for normal distribution by Shapiro-Wilks tests and were then analysed by appropriate statistical tests. Statistically significant differences were determined using Students’ t tests, or Mann Whitney U tests for non-parametric data. Where groups of data had unequal variance, a Welch’s t test was used. All statistical tests were performed using Prism Software (GraphPad Software Inc., USA). Values of p<0.05 were considered significant. All experiments were performed at least twice, with at least 3 independent samples per group.

## Results

### CD200R1 is required for efficient IL-17A production by **γδ** T cells

CD200R1-deficient mice have a reduced ability to produce IL-17A in both psoriasis and cutaneous fungal infection models (Linley, 2023). Given the importance of γδ T cells for IL-17A production in these models, we measured IL-17A production by *ex vivo* CD200R1-deficient cutaneous γδ T cells (Figure 1A), demonstrating that CD200R1 is required for efficient IL-17A production by dermal γδ T cells (Figure 1B). CD200R1 suppresses cytokine production in myeloid cells, therefore we hypothesized that in the CD200R1-deficient cultures there may be an overactive cell type present which inhibits γδ T cell IL-17 production. To test this, CD200R1-deficient dorsal skin cells were fluorescently labelled and co-cultured with unlabelled WT cells before stimulation. CD200R1-deficient γδ T cells, retained an impairment in IL-17A production when co-cultured (Figure 1C), demonstrating that CD200R1-deficient cultures do not contain factors that inhibit IL-17 production by γδ T cells. Therefore, γδ T cells are impaired in IL-17A production when isolated from CD200R1-deficient skin.

**Figure 1:**
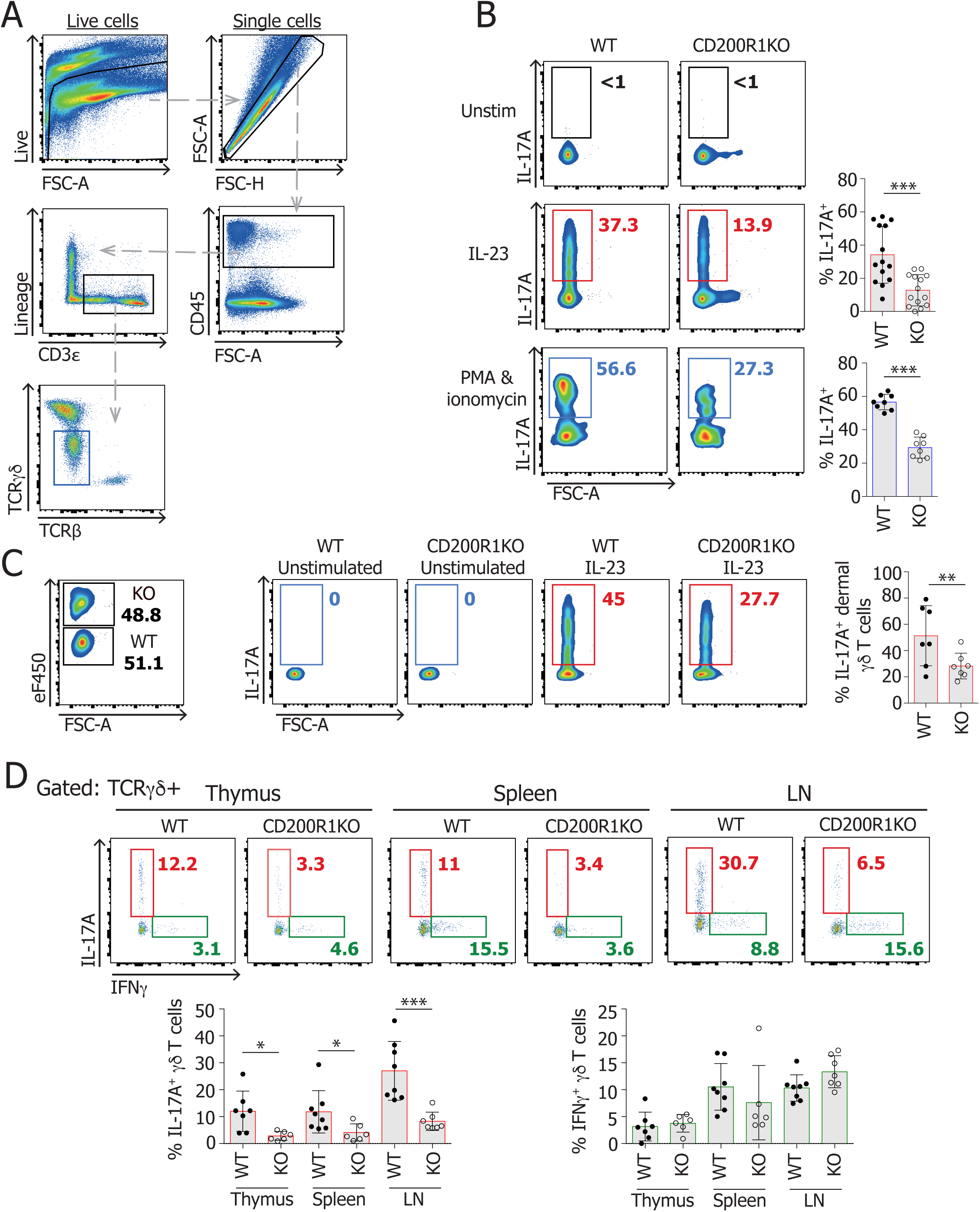
γδ T cells from CD200R1-deficient mice have an impaired ability to produce IL-17A. Cells were isolated from WT or CD200R1-deficient (KO) mouse dorsal skin then were placed in culture and stimulated with 40 ng/ml IL-23 overnight, or 50 ng/ml PMA and 500 ng/ml ionomycin for 4 hrs. IL-17A production within the TCRγδ^low^ cells was analysed by flow cytometry. **A.** Gating strategy for TCRγδ^low^ cells. **B.** IL-17A production by TCRγδ^low^ cells after stimulation. IL-23 stimulation n= 14 per group, PMA/Ionomycin stimulation n= 8 per group. **C.** Prior to culture and stimulation, CD200R1-deficient (KO) cells were labelled with a fluorescent dye, then mixed with unlabelled WT cells in a 50:50 ratio. n=7 per group. **D.** Cells were isolated from thymus, spleen and lymph nodes and were stimulated with 50 ng/ml PMA and 500 ng/ml Ionomycin for 4 hrs, before flow cytometric analysis for IL-17A and IFNγ production by γδ T cells. For thymus WT n=6, KO n=7. n=8 per group for spleen and LN. Data points indicate individual mouse data. Numbers on flow plots show percentages of cells in each gate. Data are pooled from at least two independent experiments. Students’ t-tests were used to determine statistically significant differences except where data are not normally distributed, where Mann Whitney U tests were used (**D.** IL-17 and IFNγ production in spleen and IL-17 production in LN). * signifies p<0.05, ** signifies p<0.01, *** signifies p<0.001.

To determine if CD200R1 affects γδ T cell cytokine production in a tissue specific, or γδ T cell subset-specific manner, cells from spleen, lymph node (LN) and thymus were stimulated with PMA and Ionomycin and IL-17A and IFNγ production by γδ T cells was measured by flow cytometry. WT and CD200R1-deficient γδ T cells had a similar ability to produce IFNγ, but CD200R1-deficient γδ T cells were impaired in IL-17A production in each tissue examined (Figure 1D), showing that CD200R1 promotes IL-17A production by γδ T cells, but does not affect IFNγ production.

### CD200R1 deficiency reduces **γδ**T17 cell populations and impairs their ability to produce IL-17

Reduced IL-17A production by CD200R1-deficient γδ T cells could be caused by a reduction in the IL-17-producing γδ T cell (γδT17) population, or due to these cells having a reduced ability to produce IL-17. To identify the effect of CD200R1 on T cell populations, conventional CD4^+^ αβ T cells and γδ T cells were examined in skin and lymphoid organs, showing there are not overall differences in conventional CD4^+^ αβ T cells (Figure 2A) or γδ T cells in thymus or spleen (Figure 2B). However, the overall γδ T cell population is reduced in lymph node (LN) and dermis in CD200R1-deficient mice (Figure 2B), suggesting that CD200R1 is required for optimal γδ T cell numbers in these organs.

**Figure 2:**
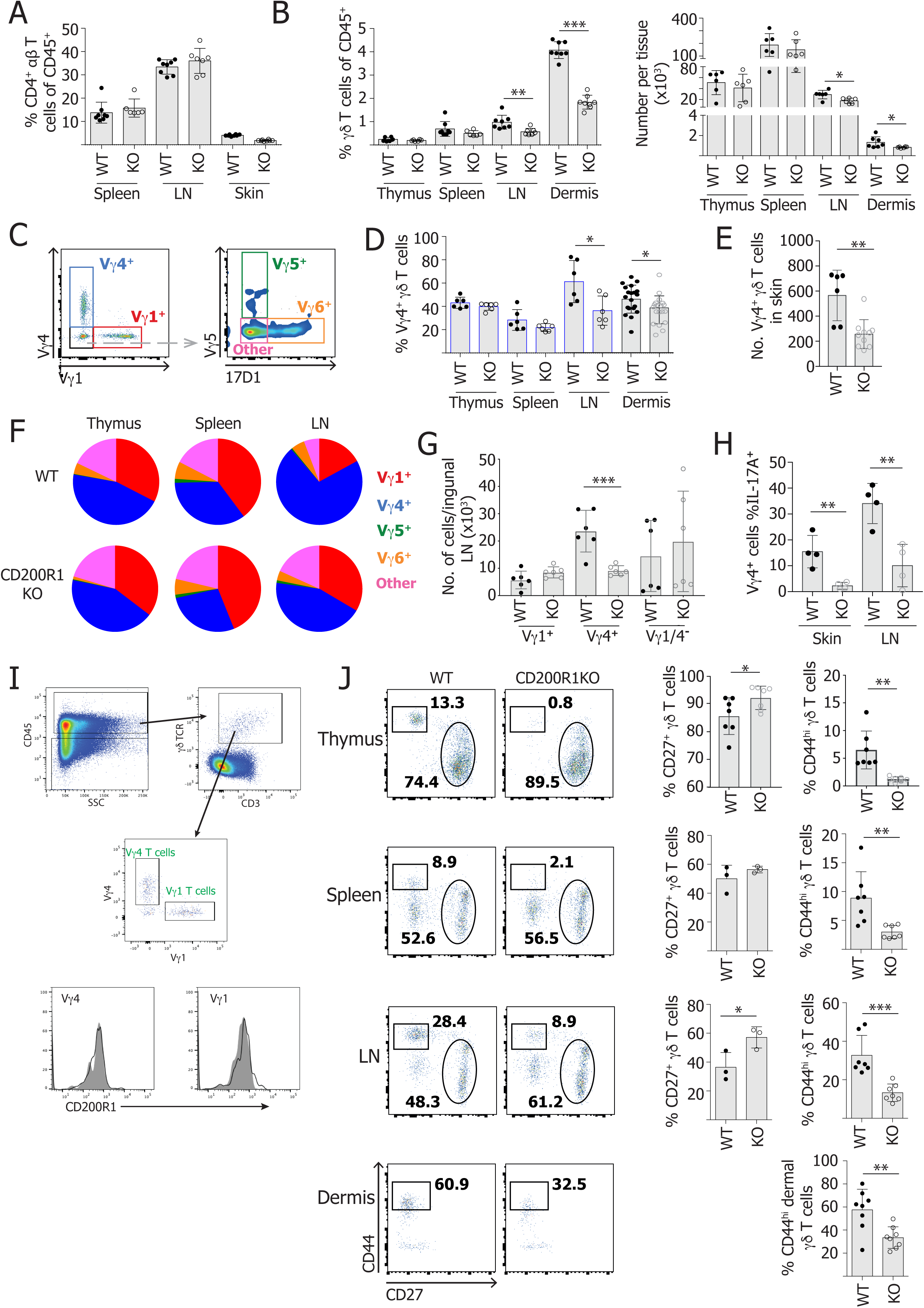
CD200R1-deficient mice have a reduced population of γδ17 T cells. Cells were isolated from thymus, spleen, LN (inguinal axillary and brachial) and skin of WT and CD200R1-deficient (KO) mice and were analysed by flow cytometry. **A.** The proportion of CD45^+^ cells that are TCRβ^+^ CD4^+^. n=8 except spleen KO n=6, LN n=7. **B.** The proportion of CD45^+^ cells that are γδ T cells in thymus, spleen and LN, or TCRγδ^low^ T cells in skin. Numbers of γδ T cells in thymus, spleen and LN, or TCRγδ^low^ T cells in skin. n=6 per group except in left panel where n=8 in spleen, LN and dermis. **C.** Representative gating for Vγ subsets in LN. **D.** The proportion of CD45^+^ cells that are Vγ4^+^ T cells in thymus, spleen, LN and skin. n=6 per group except dermis where n=20 per group. **E.** The numbers of Vγ4^+^ T cells in skin. WT n=6, KO n=10. **F.** Mean proportions of the Vγ subsets within the γδ T cell population. **G.** No. of γδ T cells within each Vγ subset in one inguinal LN. n=6 per group. **H.** Proportion of Vγ4^+^ cells producing IL-17A after PMA and ionomycin stimulation. n=4 per group. **I.** CD200R1 expression on skin Vγ4^+^ and Vγ1^+^ T cells. Filled bar shows CD200R1KO, line shows WT cells. **J.** Proportion of γδ T cells expressing high levels of CD44 (γδ17 T cells) or expressing CD27 (cells with potential for IFNγ production). Thymus n=7, spleen and LN n=3 for CD27^+^ population, n=7 for CD44^hi^. n= 8 for dermis. Data points indicate individual mouse data. At least 2 independent experiments performed. Numbers on flow plots show percentages of cells in each gate. Students’ t-tests were used to determine statistically significant differences except where data are not normally distributed, where Mann Whitney U tests were used (**G.** Vγ1/Vγ4^-^ population. **J.** CD44^hi^ data in spleen and LN). Where variances were significantly different, Welch correction was used (**G.** Vγ4^-^ population. **H.** skin data. **J.** Thymus CD44^hi^). * signifies p<0.05, ** signifies p<0.01, *** signifies p<0.001.

γδ T cells take on their effector fate during development in the thymus, when they also rearrange their TCR loci and express specific Vγ and δ TCR chains. In skin, the major IL-17A-producing subset expresses Vγ4. Indeed, in LN and dermis there is a reduction in the proportion and number of Vγ4^+^ cells in CD200R1-deficient mice (Figure 2C-G), demonstrating that CD200R1 supports this population. As well as this population being reduced in CD200R1-deficient skin and LN, the Vγ4 T cells remaining have a reduced ability to produce IL-17A in the absence of CD200R1 (Figure 2H), revealing that CD200R1 promotes IL-17 production, and is required for optimal numbers of γδT17 cells.

Previously, only low levels of CD200R1 expression were observed in dermal γδ T cells (39, 40), but specific subsets were not examined. To determine if CD200R1 expression is specific to Vγ4^+^ cells, flow cytometric staining for CD200R1 was carried out, revealing that both cutaneous Vγ4^+^ and Vγ1^+^ T cells express exceptionally low levels of CD200R1 (Figure 2I). This suggests that CD200R1 does not directly affect the activity or maintenance of Vγ4^+^ T cells once they are mature, but instead points towards either indirect effects via another cell type, and/or effects earlier during the development of these cells. The previous *in vitro* co-culture experiments (Figure 1C) show that there is not a cell type in CD200R1-deficient cultures that inhibits γδT17 activity, suggesting that CD200R1 affects γδT17 cells prior to their activation, perhaps during their development.

Although the proportion of Vγ4^+^ T cells was not changed in the thymus and spleen (Figure 2D), γδ T cells are less able to produce IL-17A in CD200R1-deficient thymus, spleen and LN (Figure 1D), suggesting that CD200R1 may indeed have a developmental impact. Therefore, γδ T cell subsets were examined in thymus, spleen, LN and dermis based on CD44^hi^ and CD27^+^ expression, corresponding largely to IL-17A and IFNγ-producers respectively (19). In all these tissues, the proportion of CD44^hi^ γδ T cells is reduced in CD200R1-deficient mice (Figure 2J), demonstrating that γδT17 cell populations are reduced in the thymus and spleen, despite the Vγ4 T cell subset not being reduced in number. Therefore, CD200R1 is required for optimal numbers of γδT17 cells including the CD44^hi^ subset in skin and lymphoid organs, and specifically the Vγ4 T cell subset in skin and LN.

### CD200R1 promotes thymocyte commitment to the **γδ**T17 cell lineage

To determine if reduced Vγ4 T cell numbers in CD200R1-deficient mouse skin and LN are due to decreased survival, or proliferation of these cells, flow cytometric analysis of anti-apoptotic Bcl-2 levels, Annexin V binding and 7AAD staining, and Ki67 expression was examined on Vγ4^+^ T cells (Supplementary Figure 1), demonstrating no effects of CD200R1 on survival, or proliferation of these cells. Therefore, together with the evidence of a reduced CD44^hi^ γδ T cell population in thymus (Figure 2J), it seems likely that there is reduced production of Vγ4^+^ T cells in CD200R1-deficient mice.

γδ T cells develop in the thymus where they take on their effector fate, before migrating to the periphery. To determine if CD200R1 affects the development of T cells generally, CD4 and CD8 expression in the thymus was investigated, showing that overall T cell development is largely normal in CD200R1-deficient mice (Figure 3A). It has previously been shown that γδ T cells in the thymus can be gated into 3 distinct pathways of development, based on the cells’ ability to generate IFNγ-producing γδ T cells (populations D and E), TCR-naïve adaptive γδ T cells (populations B and C), and γδNKT and resident γδT17 cells (populations D, F and G) (Figure 3B)(45). Gating γδ TCR-expressing thymocytes in this manner showed no significant effect of CD200R1 on adult mouse thymic γδ T cell development (Figure 3C). CD73 is an adenosine receptor which is induced in TCR γδ^+^ CD4^-^ CD8^-^ double negative (DN) cells in response to TCR engagement and marks cells committing to the γδ T cell lineage in adult thymus (46). CD200R1-deficient adult thymus has a lower proportion of cells expressing CD73 in each DN subset (Figure 3D), and a reduced overall proportion and number of CD73^+^ cells in thymic DN cells (Figure 3E), demonstrating a reduction in T cells committing to the γδ T cell lineage in the absence of CD200R1.

**Figure 3:**
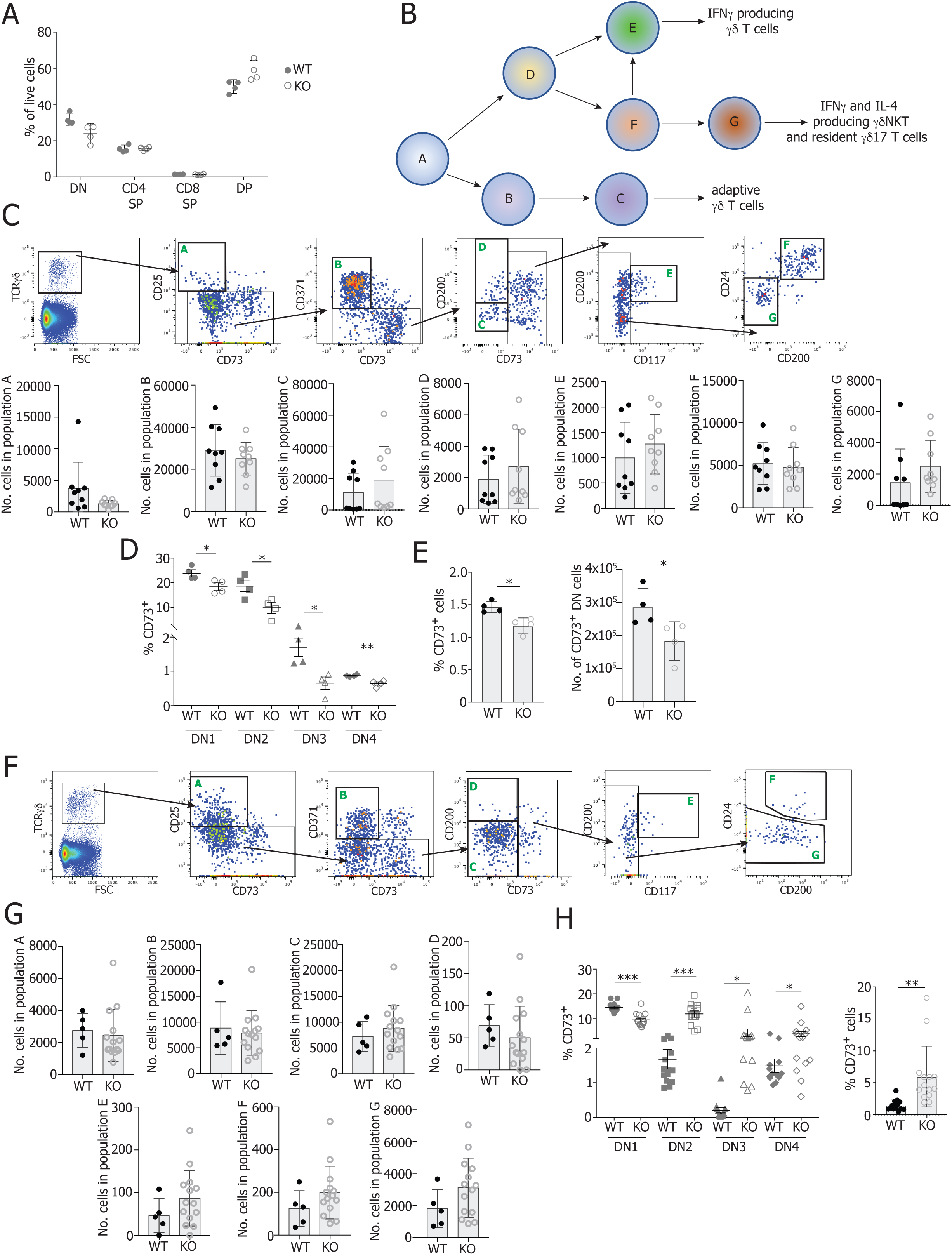
CD200R1-deficient mice have lower numbers of T cells committing to the γδ lineage. Adult, and four-day-old neonatal WT and CD200R1-deficient (KO) mice thymi were analysed for γδ T cell development and commitment to the γδ T cell lineage by flow cytometry. **A.** Thy1^+^ T cells from adults were gated into populations based on CD4 and CD8 expression. DN (double negative) CD4^-^ CD8^-^, SP (single positive), DP (double positive) CD4^+^ CD8^+^ n=4 **B.** Schematic of γδ T cell developmental pathways in adult thymus as defined by (45). **C.** Thymocytes from adult mice were gated on γδ T cells, then split into populations at different stages of the developmental pathways to generate IFNγ-producing γδ T cells (populations D and E), TCR-naïve adaptive γδ T cells (populations B and C), and γδNKT cells (populations D, F and G). Numbers of cells in each population per thymus shown. n=9 per group. **D.** Adult thymocytes were gated into DN cells (CD45^+^ Thy1^+^ CD4^-^ CD8^-^), then were split into DN1 (CD44^+^ CD25^-^), DN2 (CD44^+^ CD25^+^), DN3 (CD44^-^ CD25^+^) and DN4 (CD44^-^ CD25^+^) populations, in which CD73 expression was examined. n=4 per group. **E.** Proportion and numbers of DN thymocytes that express CD73 in adult thymus. n=4. **F.** Neonatal thymocytes gated into developmental stages. **G.** Numbers of cells in each neonatal thymocyte population. WT n =5, KO n=12. **H.** Proportion of CD73 expression on each DN subset in neonates and the proportion and number of CD73 expressing DN cells in neonatal thymus. n=14 per group. Data points indicate individual mouse data. Students’ t-tests were used to determine statistically significant differences. Where variances were significantly different, Welch correction was used (**C.** population A. **H.** DN2, DN3, DN4 and %CD73^+^ cells). * signifies p<0.05, ** signifies p<0.01, *** signifies p<0.001.

### CD200R1 promotes neonatal and foetal **γδ**T17 cell development

Although γδ T cells can develop in adult mice, the most significant generation of these cells, certainly in the absence of inflammation, occurs early in life. Therefore, the impact of CD200R1-deficiency on developing γδ T cell populations in neonates was examined, demonstrating that the numbers of cells in each developing population A-G were not affected by CD200R1 (Figure 3F-G). In addition to marking cells committing to the γδ T cell lineage in adult thymus, in neonatal thymus, CD73 is expressed at a higher level on cells committing to the IFNγ-producing, than the IL-17-producing γδ T cell lineage, reflecting stronger TCR γδ signalling (14). Examining CD73 expression in neonatal thymic γδ T cells showed an increase in CD73 expression level in CD200R1-deficient mice (Figure 3H), demonstrating an increased proportion of γδ T cells belonging to the IFNγ-producing subset relative to the IL-17-producing subset.

Vγ4^+^ T cells specifically are reduced in adult CD200R1-deficient skin and LN (Figure 2D-G). These cells develop in a wave during foetal development and seed the skin to become a resident population. Therefore, the impact of CD200R1 on foetal γδ T cell development was examined using foetal thymic organ cultures (FTOC) (gating shown in Supplementary Figure 2). Total numbers of γδ T cells were not affected by CD200R1 deficiency, but the overall number of IL-17-producing γδ T cells was reduced in the absence of CD200R1 (Figure 4A). Similarly, both the proportion and numbers of RORγt-expressing γδ T cells was reduced in CD200R1-deficient FTOC (Figure 4A) showing that CD200R1 is required for the development of γδT17 cells.

**Figure 4:**
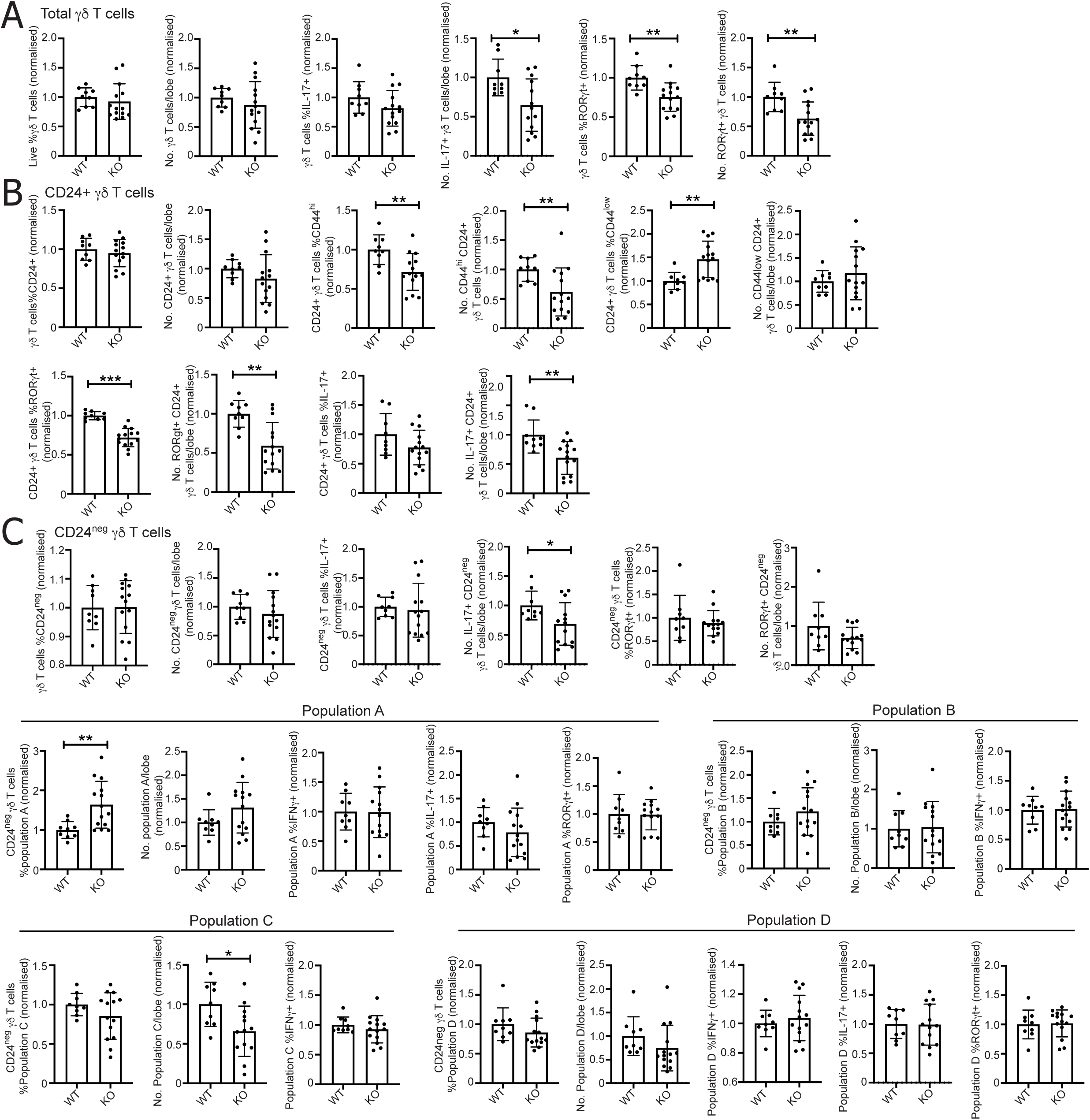
CD200R1 is required for optimal development of IL-17+ and RORγt+ γδ T cells. WT and CD200R1-deficient (KO) foetal thymic organ cultures were set up and cultured for 8 days. Cells were stimulated with PMA and Ionomycin for 3 hrs, and cell populations were examined by flow cytometry. Data are normalised to the average of the WT data for each individual experiment. Proportions and cell numbers of populations within the **A.** total γδ T cell population. **B.** CD24+ γδ T cell population. **C.** CD24^neg^ γδ T cell population. Data points indicate individual mouse data. Data are pooled from two independent experiments. n=9 for WT, n=13 for KO. Students’ t-tests were used to determine statistically significant differences with Welch’s correction for unequal variance where required (**B.** no. CD24^+^, no. CD44^hi^, %CD44^low^, no. CD44^low^, %RORγt^+^. **C.** %IL-17^+^, no. RORγt^+^, % population A, % population C). * signifies p<0.05, ** signifies p<0.01, *** signifies p<0.001.

To determine the stage of γδT17 development affected by CD200R1, cells were gated based on CD24 expression and the progenitor CD24^+^ populations were examined, showing distinct populations based on CD44 and CD45RB expression (Supplementary Figure 2). In CD200R1-deficient FTOC, the CD44^hi^ population is reduced, with a proportional increase in the CD44^low^ population which is not due to changes in the number of these cells but is due to decreases in CD44^hi^ cells (Figure 4B). The number and proportion of CD24^+^ γδ T cells expressing RORγt and the numbers producing IL-17 are reduced in the absence of CD200R1 (Figure 4B), again demonstrating a requirement for CD200R1 to support the development of γδT17 cells, particularly at this early stage of development.

The CD24^-^ γδ T cell population contains more mature developing cells, and again, the absence of CD200R1 reduced the number of IL-17 producing cells in this population (Figure 4C). These cells can be split into four populations, with population A having the potential to produce both IL-17 and IFNγ, populations B and C only containing IFNγ producers, and population D containing mainly IL-17 producers. The absence of CD200R1 led to a minor skewing towards increased population A cells and away from population C and D cells, but no other differences were observed within the individual cell populations (Figure 4C). Overall, these data demonstrate a requirement for CD200R1 for efficient acquisition of effector function (RORγt-expression and IL-17-production) in γδT17 cells, which is observed in the early stages of development.

### Effects of CD200R1 on **γδ**T17 cell development are likely either via direct effects on **γδ** T cells, or via indirect effects on thymic dendritic cells

To ascertain how mechanistically CD200R1 affects γδT17 cell acquisition of effector function, the expression of CD200R1 on different developing thymic γδ T cell populations was assessed. In adult thymus, where CD200R1-deficiency reduces the number of cells committing to the γδ T cell lineage, CD200R1 levels were negligible on the earliest developing γδ T cells (population A), and cells on the developmental pathway to generate adaptive γδ T cells (populations B and C) (Figure 5A). Significant CD200R1 levels were observed on populations D and E on the pathway to generate IFNγ-producing γδ T cells. CD200R1 levels are also significant on cells early in the pathway to become IFNγ and IL-4 producing γδNKT and long lived resident γδ17 T cells (populations D and F), but only a small proportion of cells later in the pathway (population G) express CD200R1 (Figure 5A). Whilst it is interesting that CD200R1 levels vary in different developing γδ T cell populations, the expression pattern observed does not explain why CD200R1 affects the IL-17-producing, but not the IFNγ-producing population, as CD200R1 is expressed in cells developing along both pathways. In mature dermal γδ T cells, the level of CD200R1 expressed was found to be dependent on age, with newborns expressing high levels of CD200R1 on γδ T cells, which rapidly declines to very low levels of CD200R1 expression by adulthood (Figure 5B). Therefore, CD200R1 expression was examined in developing neonatal thymic γδ T cells, demonstrating similar patterns of expression to that seen in adults, with undetectable CD200R1 expression on the earliest developing γδ T cells (population A), and cells on the developmental pathway to generate adaptive γδ T cells (populations B and C). A significant proportion of cells in populations D and E, on the pathway to become IFNγ-producing γδ T cells express CD200R1, and CD200R1 is expressed on a proportion of cells in populations D and G which are the early and late stages of the pathway to generate IFNγ and IL-4 producing γδNKT and long lived resident γδ17 T cells, but population F in neonates does not appear to express significant levels of CD200R1 (Figure 5C).

**Figure 5:**
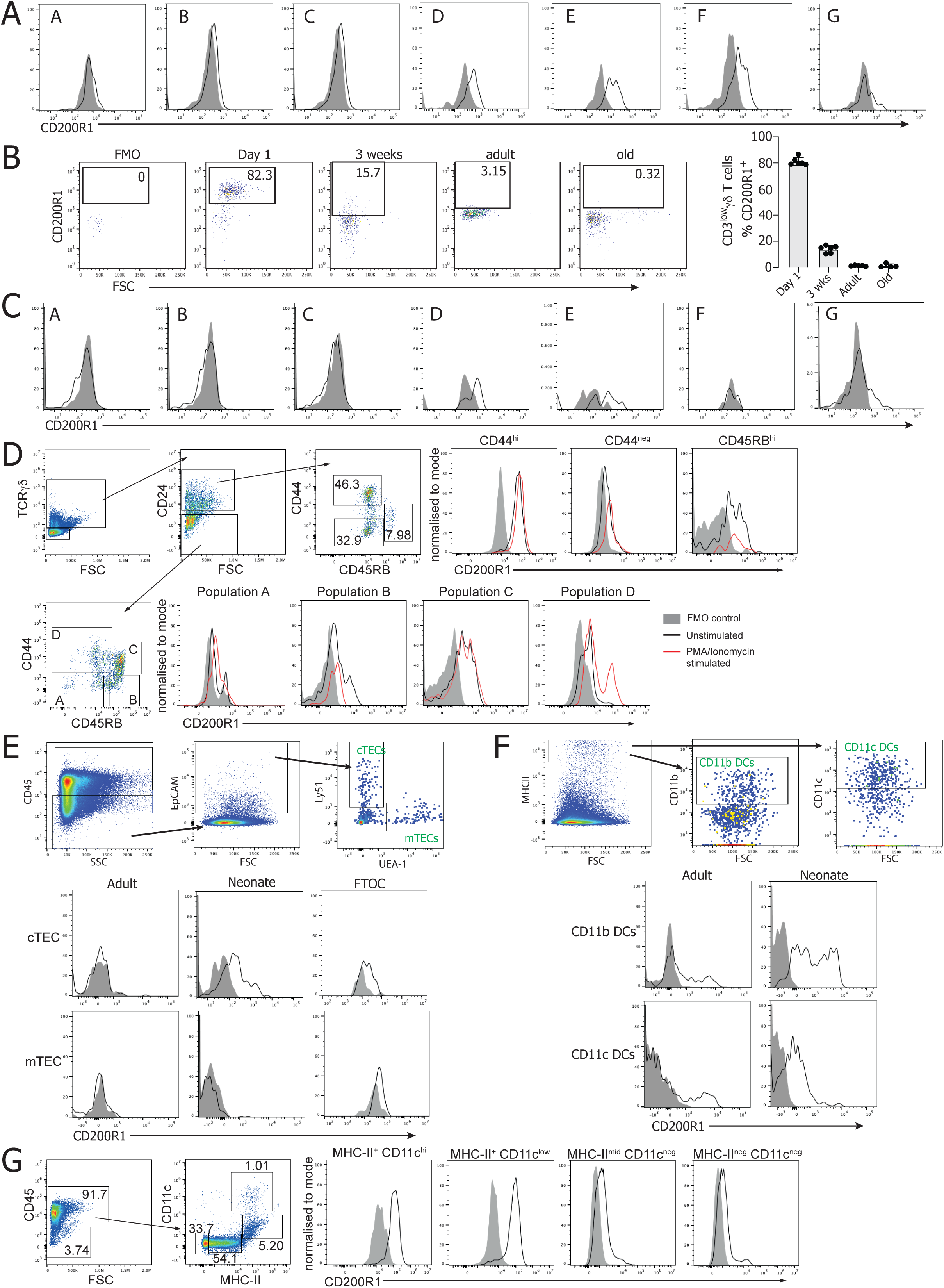
Dendritic cells, neonatal cTECs and γδ T cells express CD200R1. Flow cytometry was used to determine cell types expressing CD200R1 in skin or thymus from mice of different ages. Grey histograms depict Fluorescence minus one (FMO) controls for CD200R1 staining, black lines depict CD200R1 staining on unstimulated cells, red lines depict CD200R1 staining after a 3 hour stimulation with PMA and Ionomycin. **A.** Adult thymus was gated on γδ T cells which were split into developing subpopulations (as shown in Figure 3B-C). **B.** CD200R1 expression in cutaneous γδ T cells in one-day-old, three-week-old, adult (8-12 weeks), or old (10 months) mice. **C.** Neonatal thymus was gated on γδ T cells which were split into developing subpopulations (as shown in Figure 3B and F). **D.** CD200R1 expression on populations of developing γδ T cells in foetal thymic organ cultures (FTOC). **E.** Representative (adult mouse) flow cytometric gating on cTEC (CD45^-^ EpCAM^+^ Ly-51^+^) and mTEC (CD45^-^ EpCAM^+^ UEA-1^+^) and CD200R1 expression on adult, neonatal and foetal thymic cTEC and mTEC populations. **F.** Representative (adult mouse) flow cytometric gating on dendritic cell (DC) populations (CD45^+^ MHCII^hi^ CD11b^+^ or CD11c^+^) and CD200R1 expression on these population in adult and neonatal thymus. **G.** Thymic dendritic cell (DC) gating in foetal thymic organ cultures and CD200R1 expression on each subset. Plots shown are representative of at least 2 independent experiments, containing data from at least 3 mice.

CD200R1 expression on developing γδ T cells in FTOCs was also examined, showing that of the less mature CD24^+^ populations, the CD44^hi^ cells have particularly high CD200R1 expression, followed by the CD45RB^hi^ subset, whereas the CD44^low^ subset has only low levels of CD200R1 (Figure 5D) and expression was not significantly changed by activating the cells with PMA and Ionomycin (Figure 5D). Therefore, CD200R1 may directly affect the acquisition of effector function at this CD24^+^ CD44^hi^ stage of γδ T cell development. Examination of the CD24^neg^ population showed negligible expression of CD200R1 on population A cells (IL-17 and IFNγ producers) and very low expression on population D (IL-17 producers). Higher levels of CD200R1 expression were observed on populations B and C (IFNγ producers) (Figure 5D). Therefore, CD200R1 is not expressed highly on CD24^neg^ γδ T cell subtypes which produce IL-17, but it is expressed on some IFNγ producing subsets. Interestingly, the expression of CD200R1 was upregulated on population D (IL-17 producers) cells in response to PMA and Ionomycin stimulation (Figure 5D), suggesting that CD200R1 may be upregulated in response to activation, and thus may contribute to the development of these cells during an activation stage in thymus.

To determine if CD200R1 could mediate its effects on γδ17 T cell development indirectly via stromal cells, CD200R1 expression on thymic epithelial cells (TECs) and dendritic cells was examined by flow cytometry. CD200R1 is expressed in a proportion of cTECs in neonatal thymus and FTOCs but was not detected in adult thymus cTECs or mTECs from any life stage (Figure 5E). Therefore, CD200R1 is unlikely to be affecting γδ T cell development via effects on TECs in adults, and in early life CD200R1 could directly affect cTEC, but not mTEC activity. In DCs CD200R1 is expressed by both the CD11b+ and CD11c+ thymic DCs in adult and neonatal thymus (Figure 5F) and is expressed highly by MHC-II+ CD11c+ DCs in FTOC (Figure 5G). CD200R1 has no effect on dendritic cell or TEC numbers in adult thymus, but CD200R1 deficiency increased neonatal thymic DC numbers and decreased cTECs (Supplementary Figure 3) which may impact γδ T cell development. Therefore, CD200R1 promotes γδT17 development either via cell extrinsic mechanisms, likely involving thymic dendritic cells, cTECs or IFNγ-producing γδ T cell subsets (populations with high CD200R1 expression), or CD200R1 promotes γδT17 development in a cell intrinsic manner, likely whilst cells are expressing CD24, or during/after activation signalling.

## Discussion

γδ T cells are critical immune cells, being able to rapidly respond in both innate and adaptive manners, producing large amounts of IFNγ or IL-17. Here we show that mice deficient for CD200R1 have fewer IL-17-producing γδ T cells due to a requirement for CD200R1 for efficient development of γδ17 T cells (Figure 1B,D). This is not due to a complete block in the development of γδ17 T cells but is due to a deficiency in the effective acquisition of effector function in CD200R1-deficient mice. This is seen in spleen and thymus where the population of Vγ4^+^ cells is not reduced (Figure 2D), but where the CD44^hi^ γδ T cell population is reduced in the absence of CD200R1 (Figure 2J). This suggests that TCR rearrangement has occurred effectively, but the cells have not been able to take on their effector phenotype, involving upregulation of CD44 and RORγt, and ability to produce IL-17. This is also demonstrated in FTOC where the major differences were in RORγt^+^ and IL-17-producing cells, with numbers of the main CD24^neg^ γδ T cell population (A-D) not significantly affected by CD200R1 (Figure 4C).

An interesting increase in CD200R1 expression was observed on activation of population D (CD24^-^ CD44^hi^) γδ T cells in FTOC. The relevance of this upregulation is not known, but activation of mature adult γδ T cells did not result in up-regulation of CD200R1 (data not shown), suggesting that this may be developmental-stage, and population specific, and therefore may be important. However, it remains unknown if there are endogenous activation signals *in vivo* during development which would also be capable of inducing this peak in CD200R1 expression.

The mechanism by which CD200R1 promotes γδ17 T cell development it likely to be via a cell intrinsic mechanism, perhaps via suppression of TCR signalling, or via effects on other signalling pathways. CD200R1 is part of the immunoglobulin superfamily and is an ‘immune checkpoint’ pathway being therapeutically blocked in cancer to promote immune responses and allow the immune system to target the tumour (47). However, the evidence for CD200R1 specifically affecting T cells or TCR signalling is weak. Cytotoxic T cell function is inhibited by CD200R1 signalling (48), and blocking CD200R1 signalling promotes TCR-driven CD4^+^ T cell proliferation in Lupus patients, but there was no effect on healthy T cells (49). Therefore, a direct effect of CD200R1 on T cell activation or TCR signalling may be likely but remains to be proven.

CD200R1 may alternatively affect γδ17 T cell development via indirect effects involving other cell types. This would likely be via thymic dendritic cells which have the highest expression levels of CD200R1 however, a clear role for these cells in γδ T cell development is yet to be defined. Alternatively, CD200R1 may affect γδ T cell development via effects on cTECs which also express CD200R1, at least in early life (Figure 5E). cTECs are crucial for Vγ4 T cell activity (35) and support αβ T cell development by providing Notch signals and IL-7 (34), which could also be important for supporting γδ17 T cell development. Notably, cTEC numbers are reduced in the absence of CD200R1 (Supplementary Figure 3B), supporting this as a possible mechanism.

Together this demonstrates an important role for CD200R1 in driving early life γδ17 T cell acquisition of effector function specifically, without affecting the development of IFNγ-producing subsets. The precise mechanism by which CD200R1 drives γδ17 T cell development, and what regulates CD200R1 expression remains to be identified and it will also be important to determine if CD200R1 plays a similar role in human γδ17 T cell development however, this research is technically difficult to undertake. Understanding the development of γδ17 T cells and their acquisition of effector function is important as these cells have crucial functions in immune responses, tumour surveillance and wound healing, but they also contribute to inflammatory disease. Therefore, a better understanding the factors involved in the development and function of these important cells has the potential to improve multiple areas of health.

## Acknowledgements

This work was funded by a pre-competitive, open innovation award to the Manchester Collaborative Centre for Inflammation Research, at the University of Manchester, AstraZeneca and GSK, and a Wellcome Trust and Royal Society, Sir Henry Dale Fellowship to AS (109375/Z/15/Z). We acknowledge assistance from Gareth Howell, and the use of the University of Manchester and Lancaster University Flow Cytometry, Physiological Services Unit and Biological Services facilities. Joanne Konkel for advice and Adrian Hayday for kindly providing the 17D3 antibody against Vγ6. Dan Pennington and Nital Sumaria for invaluable guidance with FTOCs.

## References

1. Hayday AC. Gammadelta T cells and the lymphoid stress-surveillance response. Immunity. 2009;31(2):184–96.

2. Hayday A, Tigelaar R. Immunoregulation in the tissues by gammadelta T cells. Nat Rev Immunol. 2003;3(3):233–42.

3. Jameson J, Ugarte K, Chen N, Yachi P, Fuchs E, Boismenu R, et al. A role for skin gammadelta T cells in wound repair. Science. 2002;296(5568):747–9.

4. Girardi M, Oppenheim DE, Steele CR, Lewis JM, Glusac E, Filler R, et al. Regulation of cutaneous malignancy by gammadelta T cells. Science. 2001;294(5542):605–9.

5. Ribot JC, Lopes N, Silva-Santos B. gammadelta T cells in tissue physiology and surveillance. Nat Rev Immunol. 2020.

6. Bernal-Alferes B, Gomez-Mosqueira R, Ortega-Tapia GT, Burgos-Vargas R, Garcia-Latorre E, Dominguez-Lopez ML, et al. The role of gammadelta T cells in the immunopathogenesis of inflammatory diseases: from basic biology to therapeutic targeting. J Leukoc Biol. 2023;114(6):557–70.

7. Herrmann T, Karunakaran MM, Fichtner AS. A glance over the fence: Using phylogeny and species comparison for a better understanding of antigen recognition by human gammadelta T-cells. Immunol Rev. 2020;298(1):218–36.

8. Heilig JS, Tonegawa S. Diversity of murine gamma genes and expression in fetal and adult T lymphocytes. Nature. 1986;322(6082):836–40.

9. Prinz I, Silva-Santos B, Pennington DJ. Functional development of gammadelta T cells. Eur J Immunol. 2013;43(8):1988–94.

10. Bonneville M, Itohara S, Krecko EG, Mombaerts P, Ishida I, Katsuki M, et al. Transgenic mice demonstrate that epithelial homing of gamma/delta T cells is determined by cell lineages independent of T cell receptor specificity. J Exp Med. 1990;171(4):1015–26.

11. Haas JD, Ravens S, Duber S, Sandrock I, Oberdorfer L, Kashani E, et al. Development of interleukin-17-producing gammadelta T cells is restricted to a functional embryonic wave. Immunity. 2012;37(1):48–59.

12. Xiong N, Baker JE, Kang C, Raulet DH. The genomic arrangement of T cell receptor variable genes is a determinant of the developmental rearrangement pattern. Proc Natl Acad Sci U S A. 2004;101(1):260–5.

13. Carding SR, Egan PJ. Gammadelta T cells: functional plasticity and heterogeneity. Nat Rev Immunol. 2002;2(5):336–45.

14. Sumaria N, Grandjean CL, Silva-Santos B, Pennington DJ. Strong TCRgammadelta Signaling Prohibits Thymic Development of IL-17A-Secreting gammadelta T Cells. Cell Rep. 2017;19(12):2469–76.

15. Papotto PH, Goncalves-Sousa N, Schmolka N, Iseppon A, Mensurado S, Stockinger B, et al. IL-23 drives differentiation of peripheral gammadelta17 T cells from adult bone marrow-derived precursors. EMBO Rep. 2017;18(11):1957–67.

16. Lombes A, Durand A, Charvet C, Riviere M, Bonilla N, Auffray C, et al. Adaptive Immune-like gamma/delta T Lymphocytes Share Many Common Features with Their alpha/beta T Cell Counterparts. J Immunol. 2015;195(4):1449–58.

17. Jensen KD, Su X, Shin S, Li L, Youssef S, Yamasaki S, et al. Thymic selection determines gammadelta T cell effector fate: antigen-naive cells make interleukin-17 and antigen-experienced cells make interferon gamma. Immunity. 2008;29(1):90–100.

18. Turchinovich G, Hayday AC. Skint-1 identifies a common molecular mechanism for the development of interferon-gamma-secreting versus interleukin-17-secreting gammadelta T cells. Immunity. 2011;35(1):59–68.

19. Ribot JC, deBarros A, Pang DJ, Neves JF, Peperzak V, Roberts SJ, et al. CD27 is a thymic determinant of the balance between interferon-gamma- and interleukin 17-producing gammadelta T cell subsets. Nat Immunol. 2009;10(4):427–36.

20. Fahl SP, Coffey F, Kain L, Zarin P, Dunbrack RL, Jr., Teyton L, et al. Role of a selecting ligand in shaping the murine gammadelta-TCR repertoire. Proc Natl Acad Sci U S A. 2018;115(8):1889–94.

21. Sumaria N, Martin S, Pennington DJ. Constrained TCRgammadelta-associated Syk activity engages PI3K to facilitate thymic development of IL-17A-secreting gammadelta T cells. Sci Signal. 2021;14(692).

22. Laird RM, Laky K, Hayes SM. Unexpected role for the B cell-specific Src family kinase B lymphoid kinase in the development of IL-17-producing gammadelta T cells. J Immunol. 2010;185(11):6518–27.

23. Zuberbuehler MK, Parker ME, Wheaton JD, Espinosa JR, Salzler HR, Park E, et al. The transcription factor c-Maf is essential for the commitment of IL-17-producing gammadelta T cells. Nat Immunol. 2019;20(1):73–85.

24. He YW, Malek TR. Interleukin-7 receptor alpha is essential for the development of gamma delta + T cells, but not natural killer cells. J Exp Med. 1996;184(1):289–93.

25. Maki K, Sunaga S, Ikuta K. The V-J recombination of T cell receptor-gamma genes is blocked in interleukin-7 receptor-deficient mice. J Exp Med. 1996;184(6):2423–7.

26. Ye SK, Agata Y, Lee HC, Kurooka H, Kitamura T, Shimizu A, et al. The IL-7 receptor controls the accessibility of the TCRgamma locus by Stat5 and histone acetylation. Immunity. 2001;15(5):813–23.

27. Schlissel MS, Durum SD, Muegge K. The interleukin 7 receptor is required for T cell receptor gamma locus accessibility to the V(D)J recombinase. J Exp Med. 2000;191(6):1045–50.

28. Agata Y, Katakai T, Ye SK, Sugai M, Gonda H, Honjo T, et al. Histone acetylation determines the developmentally regulated accessibility for T cell receptor gamma gene recombination. J Exp Med. 2001;193(7):873–80.

29. Michel ML, Pang DJ, Haque SF, Potocnik AJ, Pennington DJ, Hayday AC. Interleukin 7 (IL-7) selectively promotes mouse and human IL-17-producing gammadelta cells. Proc Natl Acad Sci U S A. 2012;109(43):17549–54.

30. Baccala R, Witherden D, Gonzalez-Quintial R, Dummer W, Surh CD, Havran WL, et al. Gamma delta T cell homeostasis is controlled by IL-7 and IL-15 together with subset-specific factors. J Immunol. 2005;174(8):4606–12.

31. Nakamura M, Shibata K, Hatano S, Sato T, Ohkawa Y, Yamada H, et al. A genome-wide analysis identifies a notch-RBP-Jkappa-IL-7Ralpha axis that controls IL-17-producing gammadelta T cell homeostasis in mice. J Immunol. 2015;194(1):243–51.

32. Powolny-Budnicka I, Riemann M, Tanzer S, Schmid RM, Hehlgans T, Weih F. RelA and RelB transcription factors in distinct thymocyte populations control lymphotoxin-dependent interleukin-17 production in gammadelta T cells. Immunity. 2011;34(3):364–74.

33. Do JS, Fink PJ, Li L, Spolski R, Robinson J, Leonard WJ, et al. Cutting edge: spontaneous development of IL-17-producing gamma delta T cells in the thymus occurs via a TGF-beta 1-dependent mechanism. J Immunol. 2010;184(4):1675–9.

34. Abramson J, Anderson G. Thymic Epithelial Cells. Annu Rev Immunol. 2017;35:85–118.

35. Nitta T, Muro R, Shimizu Y, Nitta S, Oda H, Ohte Y, et al. The thymic cortical epithelium determines the TCR repertoire of IL-17-producing gammadeltaT cells. EMBO Rep. 2015;16(5):638–53.

36. Roberts NA, White AJ, Jenkinson WE, Turchinovich G, Nakamura K, Withers DR, et al. Rank signaling links the development of invariant gammadelta T cell progenitors and Aire(+) medullary epithelium. Immunity. 2012;36(3):427–37.

37. Cherwinski HM, Murphy CA, Joyce BL, Bigler ME, Song YS, Zurawski SM, et al. The CD200 receptor is a novel and potent regulator of murine and human mast cell function. J Immunol. 2005;174(3):1348–56.

38. Jenmalm MC, Cherwinski H, Bowman EP, Phillips JH, Sedgwick JD. Regulation of myeloid cell function through the CD200 receptor. J Immunol. 2006;176(1):191–9.

39. Linley H, Jaigirdar S, Mohamed K, Griffiths CEM, Saunders A. Reduced cutaneous CD200:CD200R1 signaling in psoriasis enhances neutrophil recruitment to skin. Immun Inflamm Dis. 2022;10(7):e648.

40. Linley H, Ogden A, Jaigirdar S, Buckingham L, Cox J, Priestley M, et al. CD200R1 promotes interleukin-17 production by group 3 innate lymphoid cells by enhancing signal transducer and activator of transcription 3 activation. Mucosal Immunol. 2023;16(2):167–79.

41. Cai Y, Shen X, Ding C, Qi C, Li K, Li X, et al. Pivotal role of dermal IL-17-producing gammadelta T cells in skin inflammation. Immunity. 2011;35(4):596–610.

42. Pantelyushin S, Haak S, Ingold B, Kulig P, Heppner FL, Navarini AA, et al. Rorgammat+ innate lymphocytes and gammadelta T cells initiate psoriasiform plaque formation in mice. J Clin Invest. 2012;122(6):2252–6.

43. Kashem SW, Riedl MS, Yao C, Honda CN, Vulchanova L, Kaplan DH. Nociceptive Sensory Fibers Drive Interleukin-23 Production from CD301b+ Dermal Dendritic Cells and Drive Protective Cutaneous Immunity. Immunity. 2015;43(3):515–26.

44. Boudakov I, Liu J, Fan N, Gulay P, Wong K, Gorczynski RM. Mice lacking CD200R1 show absence of suppression of lipopolysaccharide-induced tumor necrosis factor-alpha and mixed leukocyte culture responses by CD200. Transplantation. 2007;84(2):251–7.

45. Buus TB, Odum N, Geisler C, Lauritsen JPH. Three distinct developmental pathways for adaptive and two IFN-gamma-producing gammadelta T subsets in adult thymus. Nat Commun. 2017;8(1):1911.

46. Coffey F, Lee SY, Buus TB, Lauritsen JP, Wong GW, Joachims ML, et al. The TCR ligand-inducible expression of CD73 marks gammadelta lineage commitment and a metastable intermediate in effector specification. J Exp Med. 2014;211(2):329–43.

47. Nip C, Wang L, Liu C. CD200/CD200R: Bidirectional Role in Cancer Progression and Immunotherapy. Biomedicines. 2023;11(12).

48. Gorczynski R, Chen Z, Kai Y, Lee L, Wong S, Marsden PA. CD200 is a ligand for all members of the CD200R family of immunoregulatory molecules. J Immunol. 2004;172(12):7744–9.

49. Li Y, Zhao LD, Tong LS, Qian SN, Ren Y, Zhang L, et al. Aberrant CD200/CD200R1 expression and function in systemic lupus erythematosus contributes to abnormal T-cell responsiveness and dendritic cell activity. Arthritis Res Ther. 2012;14(3):R123.

