## Supplementary Material for "CD200R1 is required for the development of γδ17 T cells"

**Supplementary Table 1: Antibodies used for flow cytometry**

| **Antibody specificity** | **Clone** | **Manufacturer** |
| --- | --- | --- |
| Bcl-2 | BCL/10C4 | Biolegend |
| CD3ε | 145-2C11 | eBioscience |
| CD4 | RM4-5 | Biolegend |
| CD8 | 53-6.7 | Biolegend |
| CD11b | M1/70 | eBioscience |
| CD11c | N418 | eBioscience |
| CD19 | eBio1D3 | eBioscience |
| CD24 | M1/69 | Biolegend |
| CD25 | PC61 | Biolegend |
| CD27 | LG.7F9 | eBioscience |
| CD44 | IM7 | Biolegend |
| CD45 | 30-F11 | eBioscience |
| CD45RB | C363-16A | Biolegend |
| CD73 | TY/11.8 | Biolegend |
| CD90.2 | 53-2.1 | eBioscience |
| CD117 (cKit) | 2B8 | eBioscience |
| CD127 | A7R34 | eBioscience |
| CD200 | OX90 | eBioscience |
| CD200R1 | OX110 | eBioscience |
| CD371 | 5D3 | BD Bioscience |
| EpCAM | G8.8 | eBioscience |
| F4/80 | BM8 | eBioscience |
| FcεRIα | MAR-1 | eBioscience |
| Gr1 (Ly-6G/Ly-6C) | RB6-8C5 | eBioscience |
| IFNγ | XMG1.2 | BD Bioscience |
| IL-17A | eBio17B7 | eBioscience |
| Ki67 | SolA15 | eBioscience |
| Ly-51 (CD249) | 6C3 | eBioscience |
| MHCII (IA-IE) | M5/114.15.2 | eBioscience |
| RORγt | B2D | eBioscience |
| TCRβ | H57-597 | BD Bioscience |
| TCRγδ | eBioGL3 | eBioscience |
| Ter119 | TER-119 | eBioscience |
| UEA-1 | B-1065-2 | Vector laboratories Ltd. |
| Vγ1 (Vγ1.1) | 2.11 | Biolegend |
| Vγ4 (Vγ2) | UC3-10A6 | Biolegend |
| Vγ5 (Vγ3) | 536 | Biolegend |
| Vγ6 | 17D1 | In house |

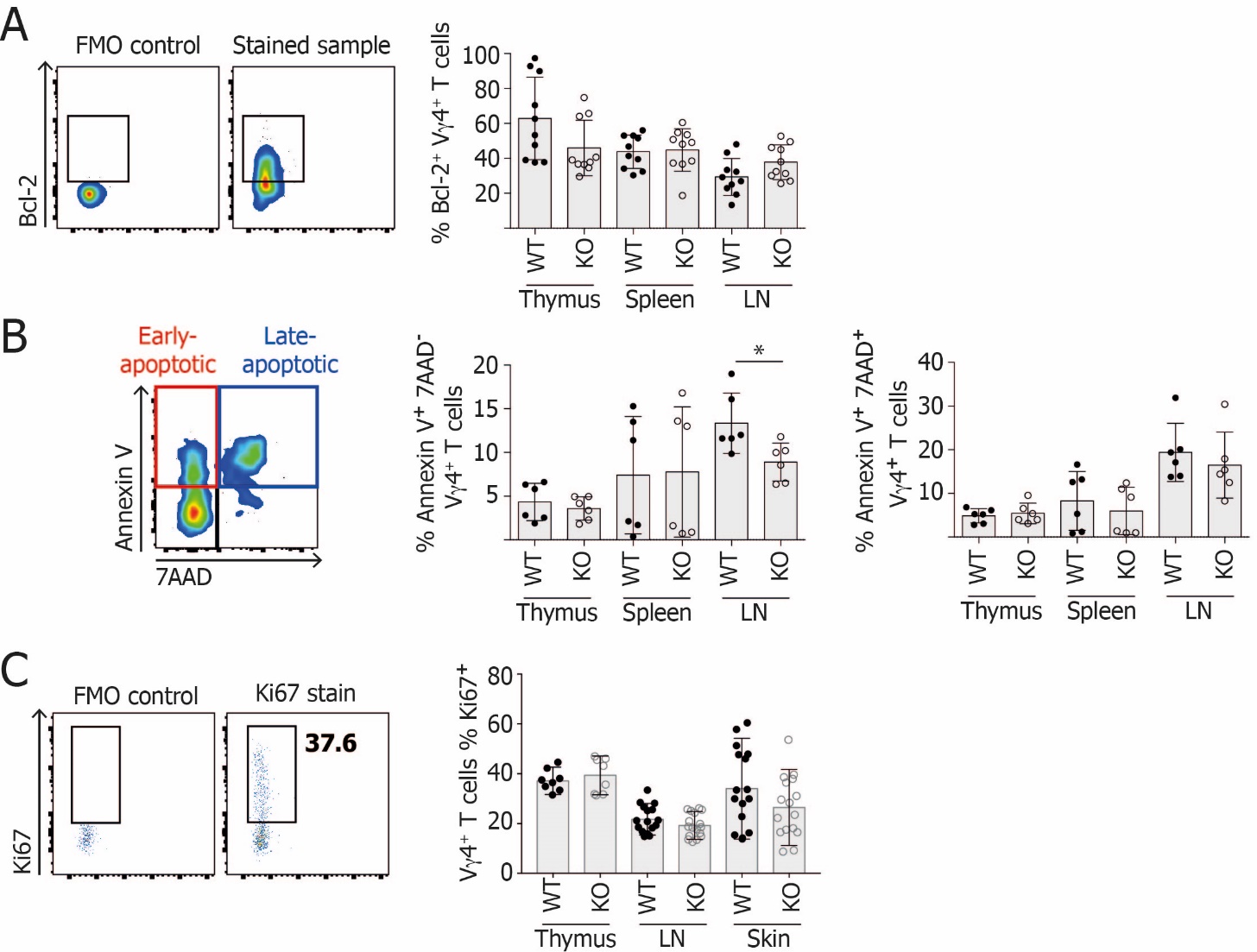

**Supplementary Figure 1: The reduction in Vγ4^+^ T cells in CD200R1-deficient mice is not due to defects in cell survival or proliferation**

Cells were isolated from thymus, spleen, LN and skin and were analysed by flow cytometry. Vγ4^+^ T cells were gated as CD45^+^ CD3^+^ Vγ4^+^ prior to gating on survival, apoptosis or proliferation markers. Representative flow plots, and bar charts with data per mouse are shown. **A.** Intracellular staining for Bcl-2. n=10. **B.** staining for Annexin V and 7AAD to examine apoptotic cells. n=6. **C.** Intracellular staining for Ki67 within the Vγ4^+^ T cell population. n=8 for thymus, n=16 for LN and skin. Students’ t-tests were used to determine statistically significant differences. Data are from at least 2 independent experiments. * signifies p<0.05, ** signifies p<0.01, *** signifies p<0.001.

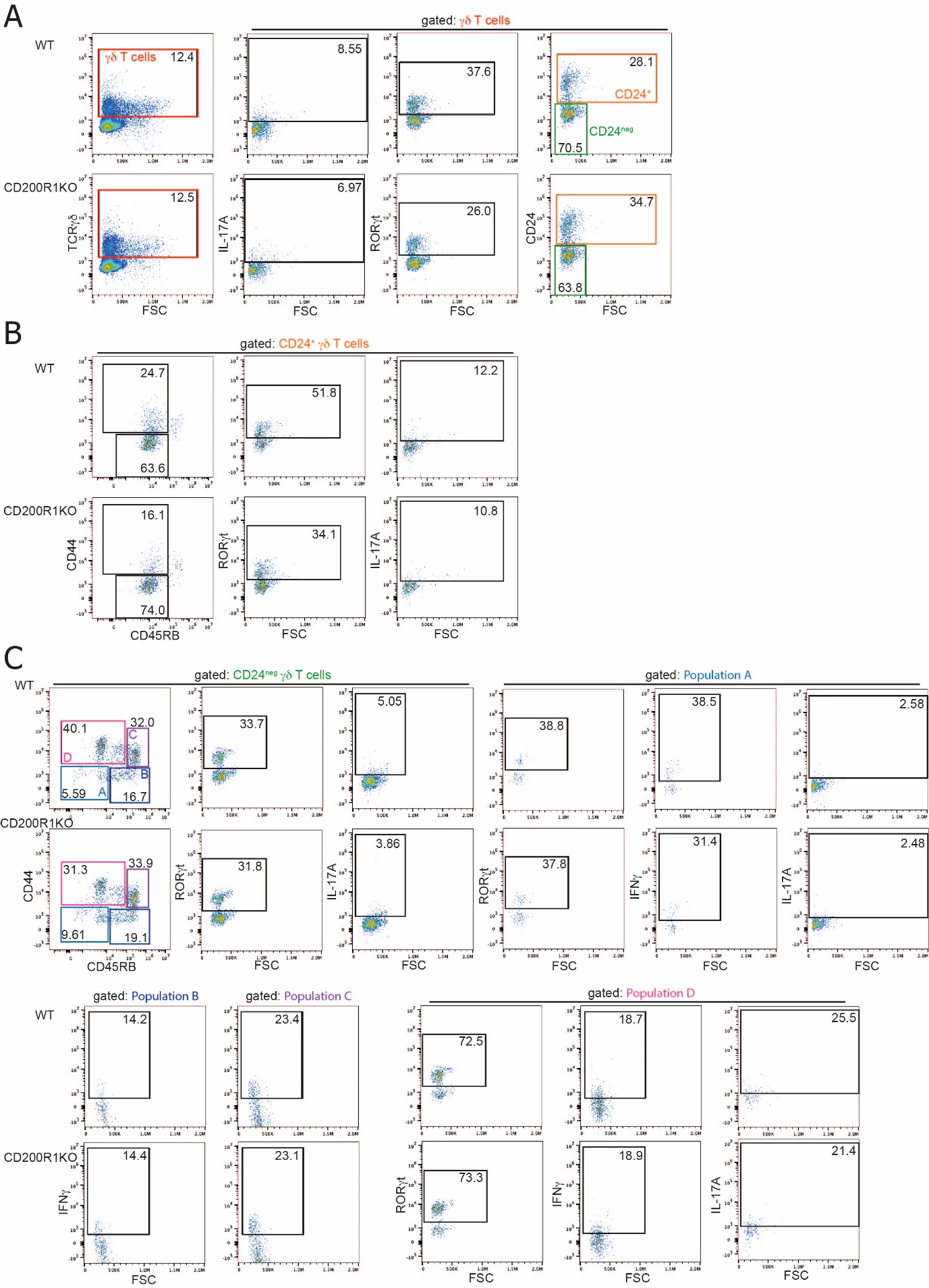

**Supplementary Figure 2: Gating strategy used to analyse developing γδ T cells in foetal thymic organ cultures**

WT and CD200R1-deficient (KO) foetal thymic lobes were cultured for 8 days. Cells were stimulated with PMA and Ionomycin for 3 hrs, and cell populations were examined by flow cytometry. Representative plots shown, with numbers indicating the proportion of cells in each gate. **A.** Gating for γδ T cells, and within this population. **B.** Gating within the CD24^+^ population of γδ T cells. **C.** Gating within the CD24^neg^ population of γδ T cells.

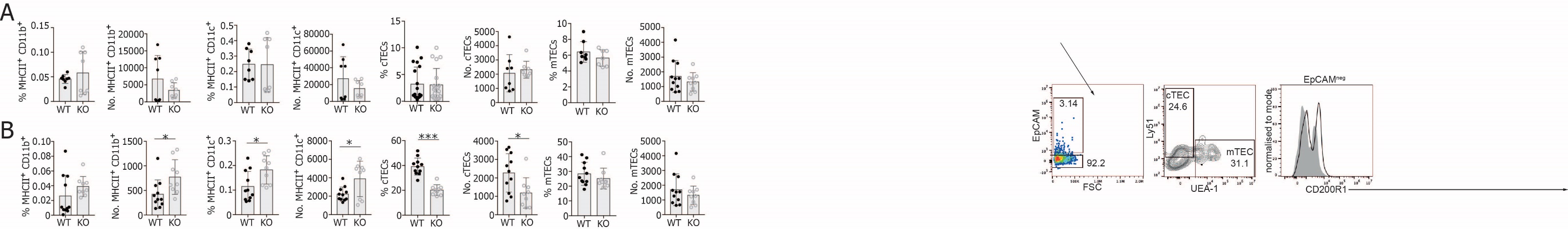

**Supplementary Figure 3: Thymic dendritic cell and cTEC populations are altered in CD200R1-deficient neonatal mice**

Adult, and four-day-old neonatal WT and CD200R1-deficient (KO) mouse thymus was analysed for dendritic cell and thymic epithelial cell (TEC) populations. Proportion and numbers of thymic dendritic cells (CD45^+^ MHCII^hi^ CD11b^+^ or CD11c^+^), cTECs (CD45^-^ EpCAM^+^ Ly-51^+^) and mTECs (CD45^-^ EpCAM^+^ UEA-1^+^) in **A.** Adult thymus. n= 8 per group, except %cTEC data, where n=19. **B.** Four-day-old neonatal thymus. WT n=11, KO n=9. Data points indicate individual mouse data. Data are pooled from two independent experiments. Students’ t-tests were used to determine statistically significant differences with Welch’s correction for unequal variance where required (**A.** % and no. MHCII^+^ CD11b^+^, no. MHCII^+^ CD11c^+^. **B.** no. MHCII^+^ CD11c^+^). * signifies p<0.05, *** signifies p<0.001.
